# A bacterial signal transduction phosphorelay in the methanogenic archaeon *Methanosarcina acetivorans*

**DOI:** 10.1101/2021.03.04.434021

**Authors:** Anne Sexauer, Nicole Frankenberg-Dinkel

## Abstract

Signal transduction via two-component systems is a powerful tool for microorganisms to respond to environmental changes. Histidine kinases originating from Bacteria are the most common signaling enzymes and are also present in Archaea, but not in all phyla. A total of 124 bacterial-type histidine kinases and/or regulators were identified in a screen of 149 Euryarchaeota genomes, but little is known about the signal transfer and molecular regulation of these systems (1).

In this work, the hybrid kinase MA4377 from the methanogenic archaeon *Methanosarcina acetivorans* was investigated (2, 3). MA4377 is a multidomain protein resembling a bacterial-type histidine kinase with two additional receiver domains at the C-terminus. Recombinant protein was employed to investigate the intra- and intermolecular phosphorelay *in vitro*. The kinase displays autophosphorylation activity of histidine residue 497. While no intramolecular phosphorelay was observed, the CheY-like receiver protein MA4376 was identified as part of the multi-component system that also seems to include the Msr-type transcription factor MA4375. This study reveals the presence and *in vitro* function of a bacterial-type hybrid histidine kinase integrated into an archaeal phosphorelay system.

**Importance:** Signal transduction enables organisms to rapidly react to changes in their surroundings. Different systems containing one-, two- or multiple-components are employed to sense and react to environmental changes. Most commonly, external perceived stimuli are converted into internal signals through protein phosphorylation. These systems are found in all domains of life but understood at different levels of complexity, the least being those from the domain of Archaea. By better elucidating the function of these early signal transduction systems, we will gain insight and understanding of the selection pressures on signal transduction pathways, and their evolution within the Archaea.

## Introduction

In order for organisms to survive, it is essential to rapidly react to environmental changes by perceiving and processing signals, thereby triggering an adequate cellular response. In bacteria, the most commonly used signal transduction system is the phosphorelay of two-components consisting of a sensory histidine kinase (HK) and a response regulator (RR) (4–8). After perceiving a signal, the kinase gets autophosphorylated at a conserved histidine residue (His1). Subsequently, the phosphoryl group is transferred to a conserved aspartate residue (Asp1) in the receiver domain (REC) of the cognate RR (Fig. 1) (4, 5, 7, 9). The phosphorylated RR then initiates the cellular response to the perceived signal, most commonly through transcriptional activation/repression. However, RR with an enzymatic function instead of DNA-binding capacity also exist and function as stand-alone receiver (REC) domains (10). In addition to this classical bacterial two-component system (TCS), multi-component systems (MCS) evolved expanding the TCS to a four-step phosphorelay (His1-Asp1-His2-Asp2) (Fig. 1). These systems consist of a hybrid histidine kinase (His1) that transfers the phosphoryl group after autophosphorylation, via a C-terminal fused receiver domain (Asp1) to a conserved histidine residue (His2) of a fused or stand-alone histidine phosphotransfer domain (HPt) that, in turn, serves as an intermediate in the phosphorelay system (11). Finally, the output RR in the MCS is phosphorylated at a conserved aspartate residue (Asp2), initiating the cellular response (4).

**Figure 1.**
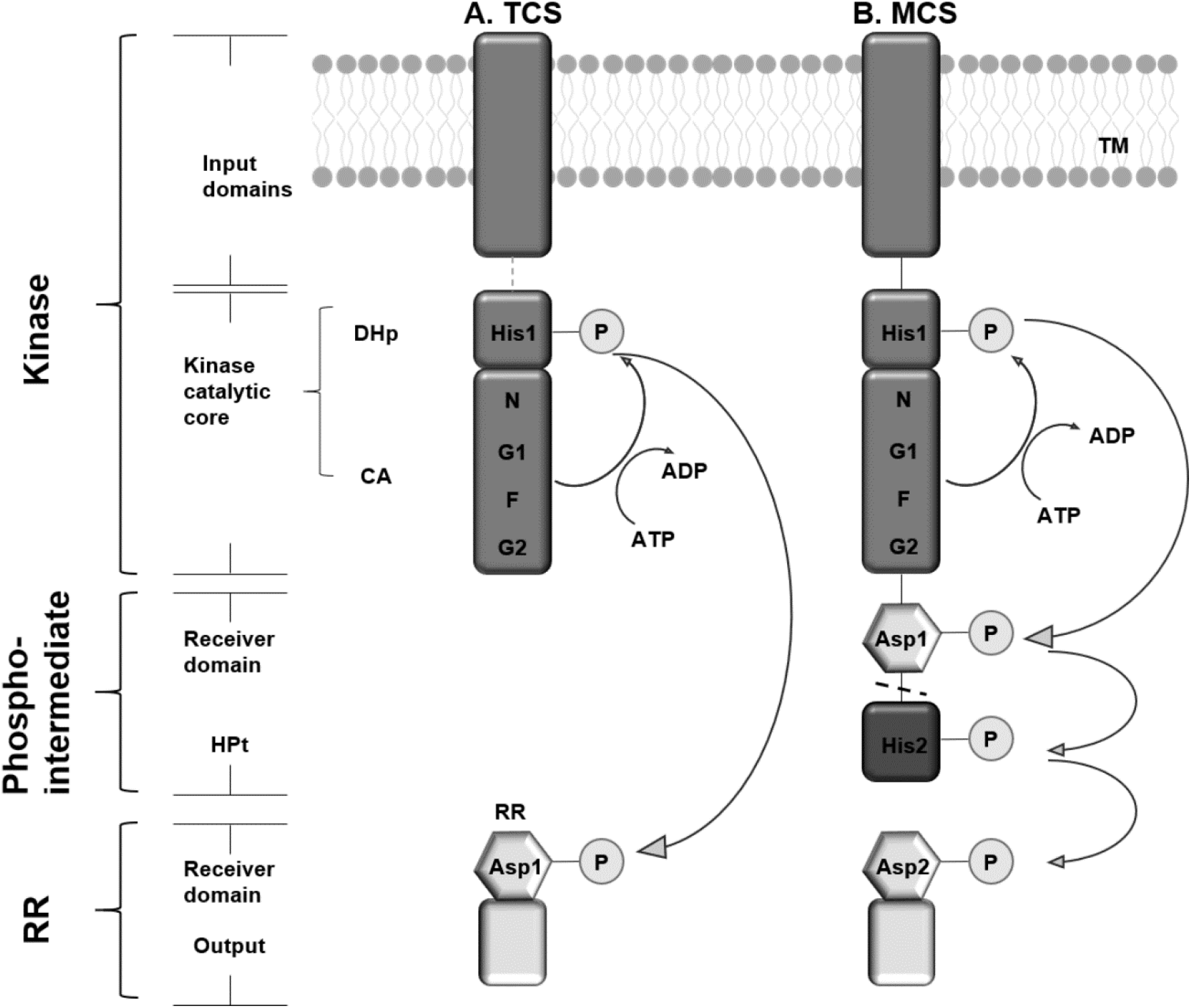
Domain organization and signal transduction of two- and multi-component systems. **A.** A classical two-component system (TCS) consists of a histidine kinase (HK) and a response regulator (RR). The HK can sense an extracellular signal, often via a membrane-anchored input domain, which is followed by the kinase catalytic core containing a DHp and a CA domain. ATP is hydrolyzed in the CA domain and the phosphoryl group is transferred to a specific histidine (His1) residue in the DHp domain. Subsequently, the phosphoryl group is transferred to a conserved aspartate residue (Asp1), located in the receiver domain of the RR, possessing an output domain. **B.** In a multi-component system (MCS), the hybrid histidine kinase is autophosphorylated at a conserved histidine residue (His1) in the DHp domain. Subsequently, the phosphoryl group is transferred to a C-terminal fused receiver domain (Asp1) and further to a histidine phosphotransfer (HPt) domain (His2) that can be either fused or stand-alone (dashed line). The next phosphate acceptor in the phosphorelay is a RR with a conserved aspartate residue (Asp2) located in the receiver domain and the following output domain. Adapted from (12).

The first TCS identified in Archaea was CheA/CheY of *Halobacterium salinarium* in 1995 (13). Since then, knowledge about signal transduction in Archaea has been steadily increasing. Genomic analysis revealed an abundance of 4000 histidine kinases and regulators in 218 archaeal genomes. None were however identified in the Cren-, Kor- and Nanoarchaeota. Screening of 149 genomes belonging to the phylum Euryarchaeota, including *M. acetivorans*, revealed that about half of the genomes contained bacterial-type histidine kinases and regulators whereas, in its subordinate class of Methanomicrobia, 3-4 % of the genes encoded a histidine kinase (1). This shows a relatively high abundance of TCS in Archaea. Thus far, only a few signaling systems have been investigated within the methanogens. Among them, the TCS LtrK/R of *Methanococcoides burtonii,* which is involved in temperature-response (14). Additionally, the TCS FilL and its corresponding regulators FilR1+2 were shown to regulate methanogenesis of *Methanosaeta harundinaceae* (15).

The methanogenic archaeon, *Methanosarcina acetivorans,* encodes 64 histidine kinases and five hybrid kinases (3, 16). Our group has previously characterized two of those sensor kinases, MsmS (MA4561) (17) and RdmS (MA0863) (18) with both resembling non-histidine kinases. Within this study, we have characterized the hybrid histidine kinase, MA4377, and its involvement in a phosphorelay system.

MA4377 possesses three sensor domains and two receiver domains fused to the kinase catalytic core. Autophosphorylation activity of a truncated version of the kinase was analyzed and the phosphorylation site of MA4377 identified. For the investigation of the four-step phosphorelay of the MCS, intra- and intermolecular phosphotransfer assays were conducted to identify signal transfer in this bacterial type MCS.

## Results

### MA4377 of *M. acetivorans* is a predicted bacterial–type hybrid kinase

MA4377 is a single open reading frame in the genome of *M. acetivorans* and annotated as a putative hybrid histidine kinase (2, 3). Upstream of the kinase gene, a methyltransferase, and an ATPase are encoded (3). Downstream, the gene for a single bacterial-type receiver domain is located, overlapping with MA4377 by 29 nucleotides, but possessing its own promoter. Adjacent to the putative receiver MA4376, but encoded on the opposite strand, genes encoding for a putative transcription factor (TF) belonging to the Msr family and an S-adenosyl-methionine (SAM)-dependent methyltransferase are positioned (Fig. 2).

**Figure 2.**
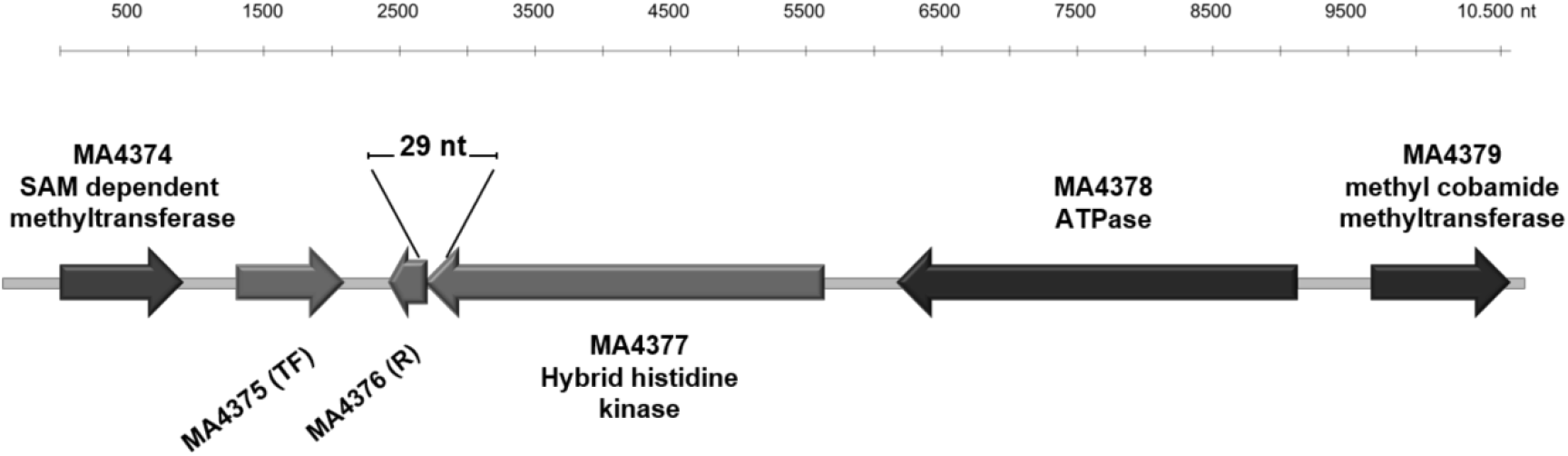
Genetic localization of MA4377 in the genome of *M. acetivorans*. Schematic representation of the genomic arrangement of MA4377, the receiver MA4376, and transcription factor (TF) MA4375 in *M. acetivorans* drawn to scale (3).

To get detailed information about the open reading frame (ORF) of MA4377, BLASTp (protein blast) analysis (NCBI; Phyre2; hhpred (MPI Bioinformatics Toolkit)) were performed to reveal the annotated domains and the similarity to other kinases supporting the characterization of MA4377. Subsequent amino acid sequence alignments (Clustal omega) using matches with the lowest E-value and the highest identity (ID) of at least 30 % but also well-characterized bacterial histidine kinases, helping to identify conserved motifs, were performed (Table S4). MA4377 encodes a predicted multidomain protein, consisting of three input domains (NCBI). The first domain represents a membrane-anchored <u>c</u>yclase/<u>h</u>istidine kinase <u>a</u>ssociated <u>s</u>ensory <u>e</u>xtracellular (CHASE4) domain (19), followed by a cytoplasmic domain present in <u>h</u>istidine kinases (HKs), <u>a</u>denylyl cyclases, <u>m</u>ethyl-accepting chemotaxis proteins (MCPs), and <u>p</u>hosphatases (HAMP) (20). C-terminal of the HAMP domain a <u>P</u>er-<u>A</u>RNT-<u>S</u>im (PAS) domain (21) as third input is succeeding. C-terminally, the kinase catalytic core, consisting of the <u>d</u>imerization and <u>h</u>istidine <u>p</u>hosphotransfer (DHp) domain and <u>c</u>atalytic and <u>A</u>TP binding (CA) domain, is located followed by two successive receivers (R1+R2) domains (Fig. 3A).

**Figure 3.**
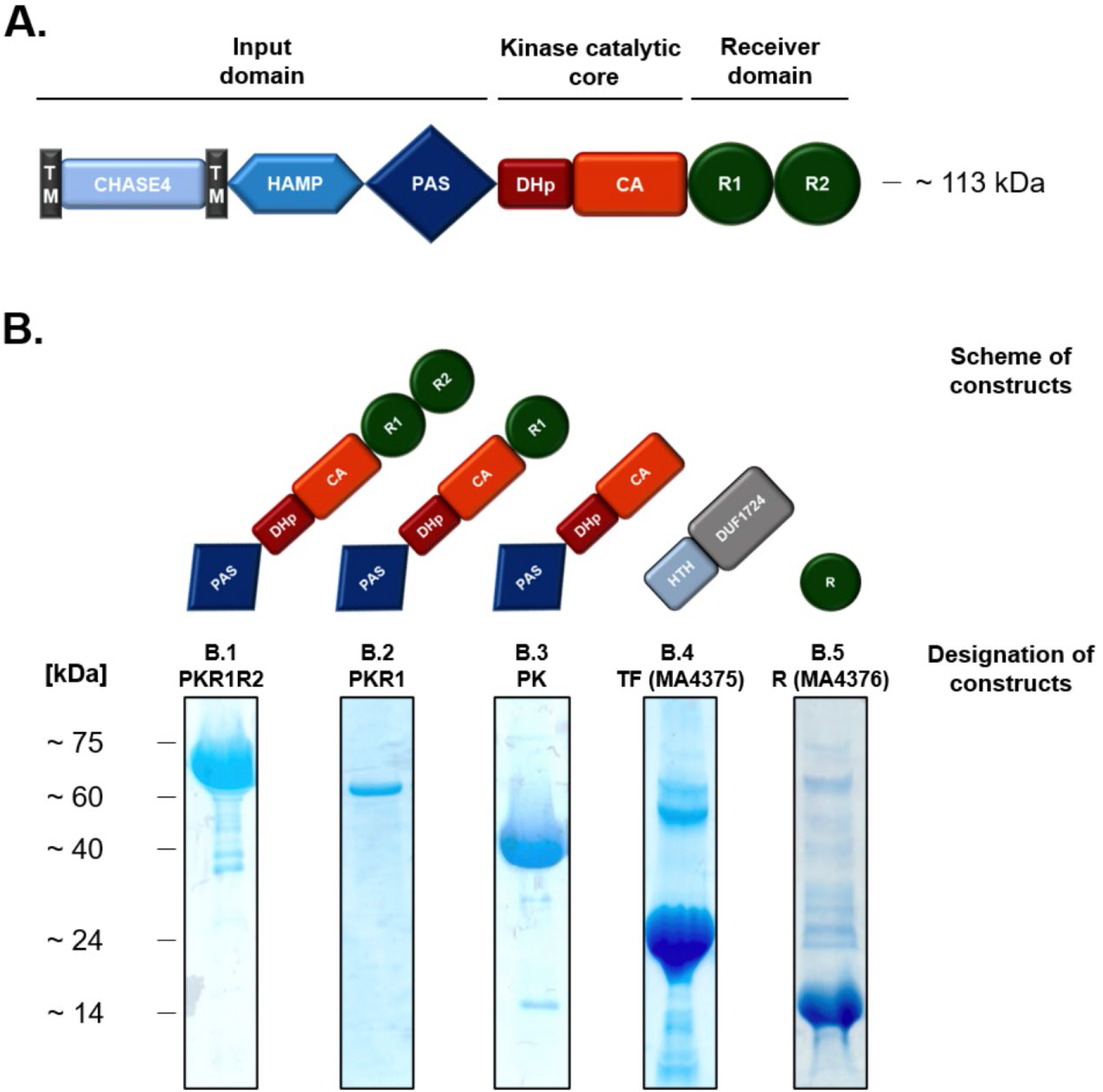
Schematic representation of the domain structure of the hybrid histidine kinase MA4377 and purification of the indicated truncated versions of MA4377. **A.** Similar annotated domains of MA4377 analyzed by tools of NCBI (BLASTp; E-value: 0.05) and hhpred (MPI Bioinformatics Toolkit) (BLASTp; E-value of 1e-3). Single domains are divided into subgroups. CHASE-, HAMP- and PAS domain (blue) are associated with input domains. DHp and CA domain (red) belong to the kinase catalytic core and R1 and R2 (green) to receiver domains. **B.** Schematic representation and SDS-PAGE of purified truncated versions used. **B.1** PKR1R2 (~75 kDa) consisting of PAS domain, the kinase catalytic core and the receiver domains R1 and R2. **B.2** PKR1 (~60 kDa) consisting of a PAS domain, the kinase catalytic core and the receiver domains R1**. B.3** PK (~40 kDa) consisting of PAS domain, the kinase catalytic core. **B.4** TF (MA4375) (~24 kDa) consisting of a helix-turn-helix (HTH) motif and a domain of unknown function 1724 (DUF1724). **B5.** R (MA4376) (~14 kDa) consisting of a receiver domain.

Based on the domain structure, connecting the kinase catalytic core with the receiver domains and the highly conserved histidine residues in the DHp domain, MA4377 resembles a bacterial-like hybrid histidine kinase.

In order to determine whether MA4377 possesses kinase activity *in vitro* and to understand its putative function in *M. acetivorans*, truncated versions of the gene were cloned for heterologous expression in *Escherichia coli*.

### The recombinant hybrid kinase MA4377 displays autokinase activity

Truncated versions of the kinase were recombinantly produced in *E. coli* BL21 (DE3) and purified via affinity chromatography to almost homogeneity (Fig. 3B). Kinases usually form dimers with the autophosphorylation occurring from one monomer to the other (22). Therefore, the oligomeric state of truncated versions of MA4377 was determined using size exclusion chromatography (Fig. S1). The truncated version of MA4377 designated PK consisting of the PAS and the kinase catalytic core (Fig. 3B) eluted as a stretched dimer from the size exclusion column. To analyze autophosphorylation activity of MA4377, the truncated version PK was employed for *in vitro* autophosphorylation assays. Autophosphorylation steadily increased over 10 min, indicating that MA4377 possesses autokinase activity (Fig. 4).

**Figure 4.**
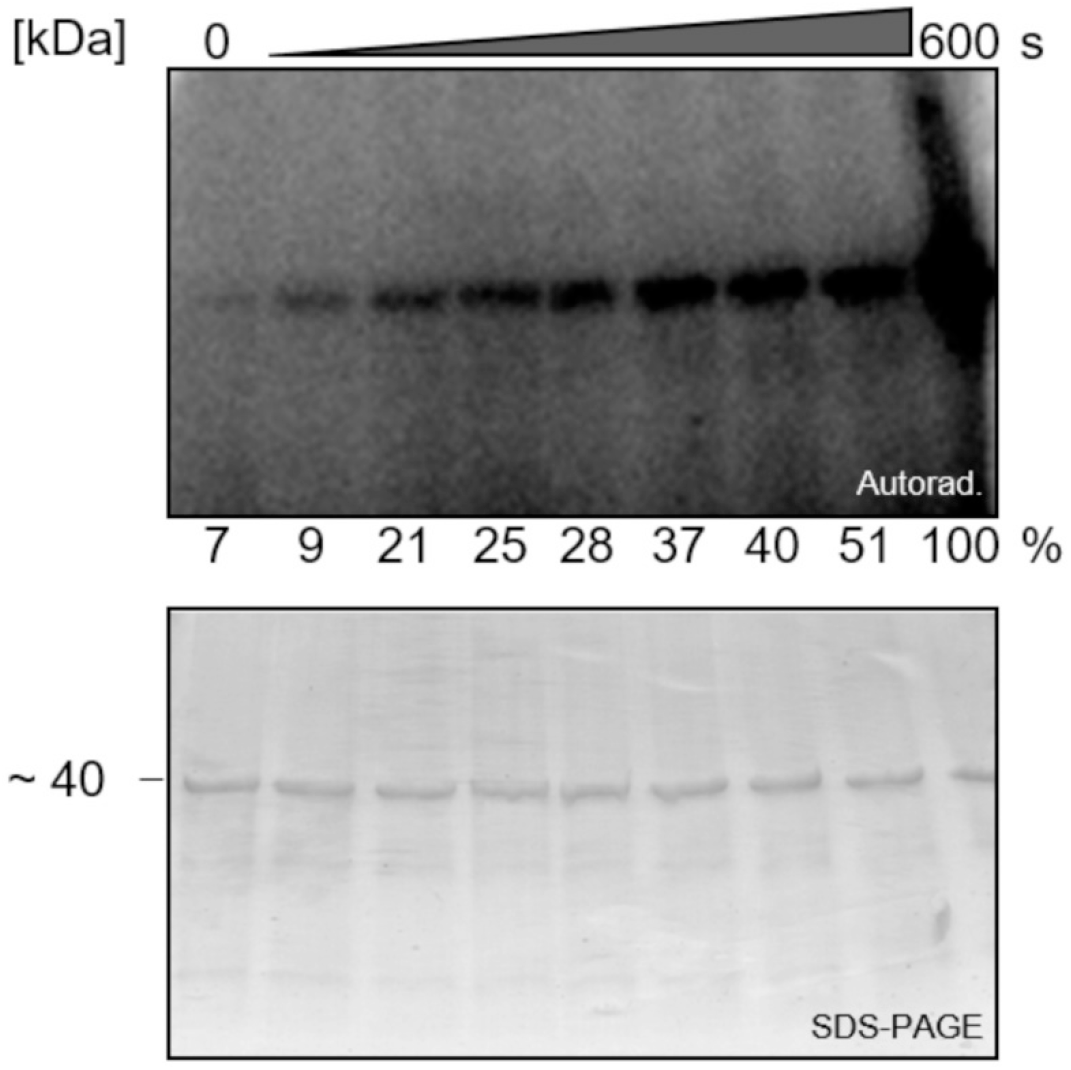
Autophosphorylation assay of the truncated version PK of MA4377.

Purified recombinant PK (10 μM) was phosphorylated with [γ-^32^P]-ATP and stopped at indicated time points using 4x SDS sample buffer. Samples (10 μl) were resolved via SDS-PAGE and subjected to autoradiography (Autorad.) or Coomassie staining to visualize the loaded protein (SDS-PAGE). The relative intensities of the phosphorylation signals are given under the autoradiogram. Values were quantified with ImageJ Color_Pixel_Counter Plugin and normalized setting 600 s sample as 100%.

### The archaeal hybrid kinase MA4377 is a histidine kinase

Signal transduction in bacteria is mostly executed by histidine kinases that get autophosphorylated on a conserved histidine residue within the so-called H-box, located in the DHp domain (Fig. 3A) (23). To investigate if MA4377 is autophosphorylated at a histidine residue, radioactively labeled kinase was subjected to acid/base treatment. Histidine residues form a phosphoramidate bond with phosphate which is acid labile and leads to a decreased radioactive signal after incubation in acid (24). MA4377 was confirmed to be a histidine kinase, as the phosphorylation was stable in alkaline solution and labile in acid (Fig. 5A). Further verification was obtained by treating the truncated versions PK and PKR1R2 (Fig. 3B1; B3) with the histidine modifying reagent diethylpyrocarbonate (DEPC) prior to the kinase assay (Fig. 5B). Upon autoradiography, no autophosphorylation signal was detected for the truncated versions, further confirming that MA4377 is a histidine kinase.

**Figure 5.**
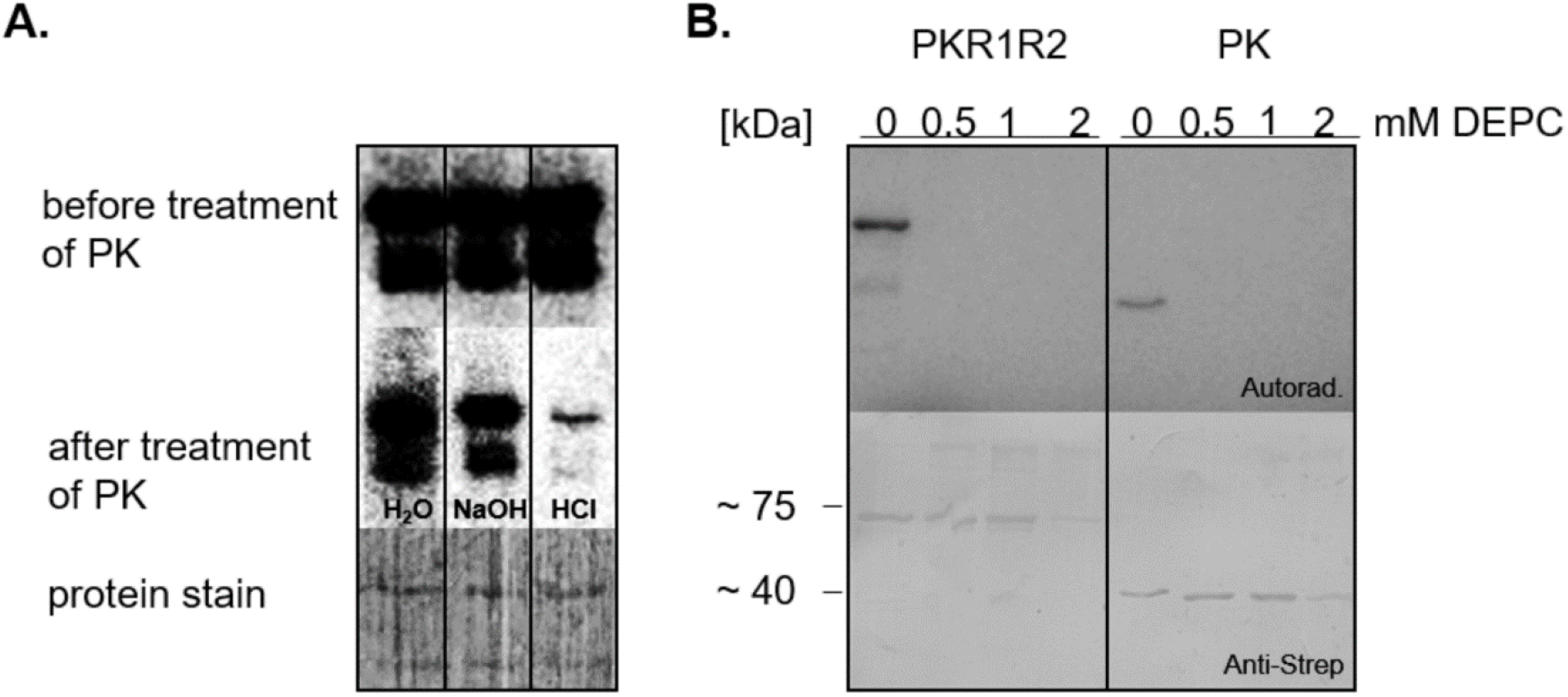
Identification of phosphorylation site. **A.** Autoradiogram of MA4377 version PK. After [γ-^32^P]-ATP radiolabeling of PK, SDS-PAGE and electroblotting, the stability of the phosphorylation was tested using acid/base treatment. As control PVDF membrane was stained with Ponceau S. **B.** Analysis of phosphorylation site by incubation 10 μM of PKR1R2 and PK with increasing concentrations 0.5, 1, 2 mM of DEPC (diethylpyrocarbonate) prior to [γ-^32^P]-ATP radiolabeling. The samples were resolved by SDS-PAGE followed by electroblotting and thereafter subjected to autoradiography showed in the upper panel as autoradiogram (Autorad.). The protein amount of the Autorad. was verified by antibody detection via Western blot (Anti-Strep).

### His_497_ is the phosphorylation site in MA4377

Amino acid sequence alignments of the DHp domain of MA4377 with kinases identified with BLASTp in *M. acetivorans* showing the lowest E-value and the highest ID of at least 30 % revealed three conserved histidine residues at positions 497, 538 and 560 (Fig. S2). To identify the specific histidine phosphorylation site, different variants were generated whereby the histidine residues were replaced by glutamine residues via site directed mutagenesis. The resulting protein variants were purified via affinity chromatography and subjected to autophosphorylation assays (Fig. 6).

**Figure 6.**
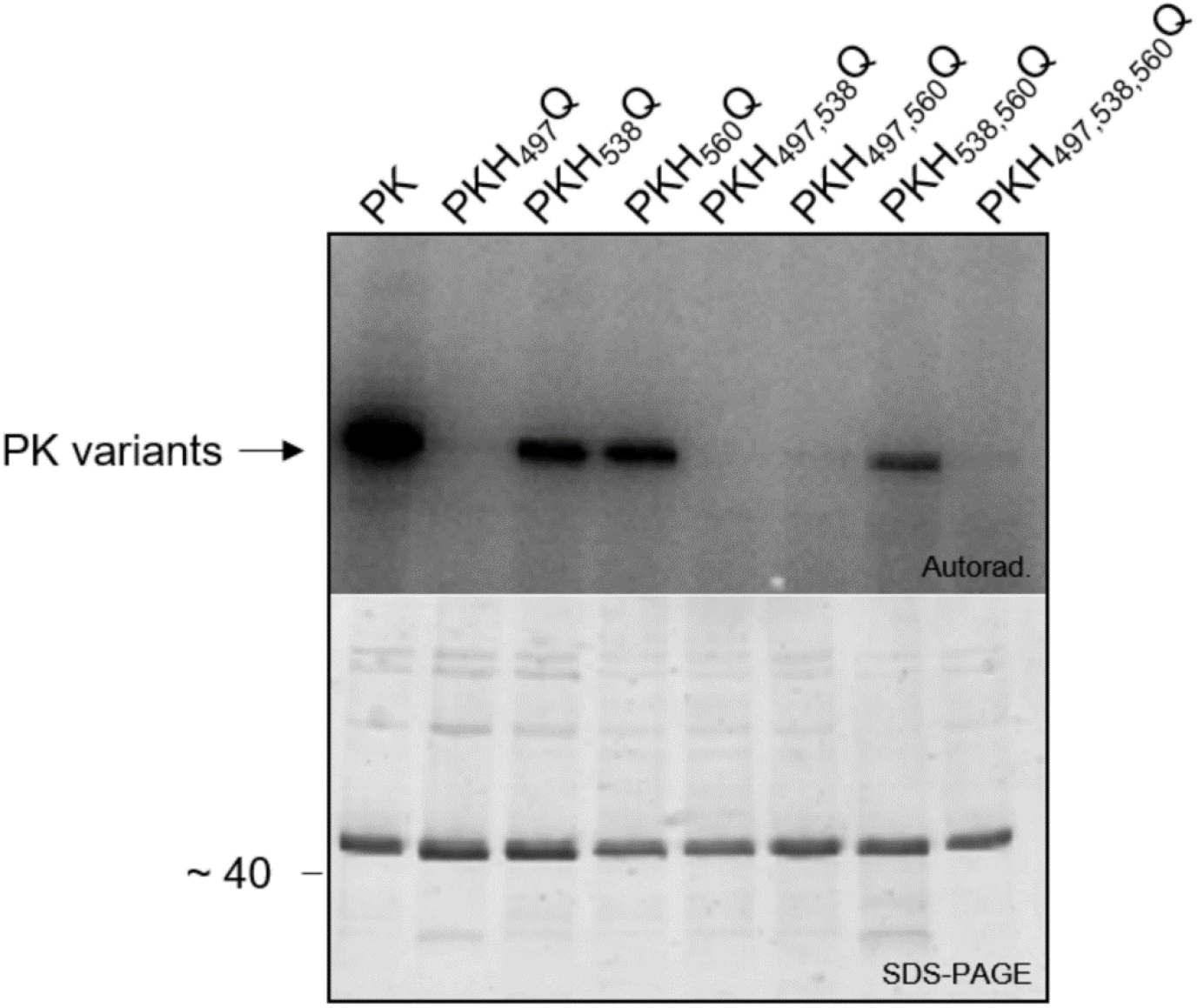
Identification of the histidine phosphorylation site. Analysis of autophosphorylation activity of the conserved histidine residue variants PKH_497_Q, PKH_538_Q, PKH_560_Q, PKH_497,538_Q, PKH_497,560_Q, PKH_538,560_Q, PKH_497,538,560_Q. [γ-^32^P]-ATP radiolabeled samples were separated by SDS-PAGE and the radioactive signals detected by PhosphoImager (Autorad.). The same SDS gel was stained with Coomassie (SDS-PAGE) to visualize the loaded protein.

Only protein variants lacking the histidine residue at position 497 showed no autophosphorylation, identifying this histidine residue as the site of phosphorylation.

### The fused receiver domains within MA4377 are not part of the phosphorelay *in vitro*

Classical bacterial signal transduction systems resemble a TCS involving a His1-Asp1 phosphotransfer from the kinase to its cognate response regulator (5, 6, 8). More complex signaling pathways are represented by hybrid histidine kinases (His1), which transfer the phosphoryl group to phosphointermediates like C-terminally fused receiver domains (Asp1) and subsequently to histidine phosphotransfer (HPt) domains (His2), until the final acceptor (Asp2) is phosphorylated, initiating an output reaction (4, 16). MA4377 possesses two additional receiver domains (R1+R2) containing conserved phosphoacceptor aspartate residues within the polypeptide chain. We therefore asked, whether the phosphoryl group from His_497_ is subsequently transferred to either of these domains. Therefore, two truncated versions PKH_497_QR1 and PKH_497_QR1D_818_NR2 both lacking the autophosphorylation site, His_497_, were generated (Fig. 7). To track the potential phosphotransfer to R1, the construct PKH_497_QR1, in combination with PK, was employed and the phosphotransfer to the R2 of MA4377 was investigated using the construct PKH_497_QR1D_818_NR2. The latter was missing the potential phosphorylation site D_818_ within receiver domain 1.

**Figure 7.**
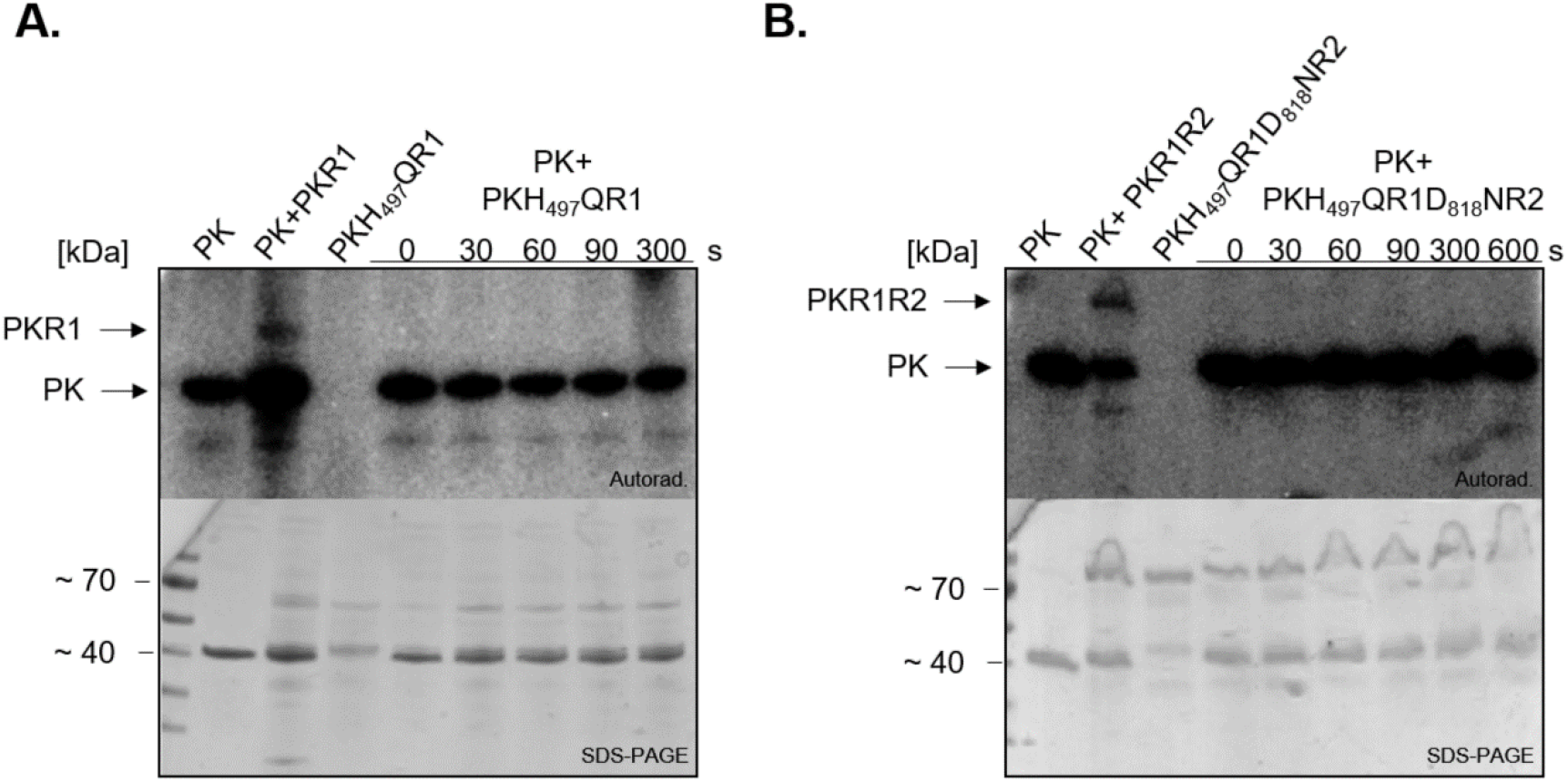
Intra-molecular phosphor transfer assay in MA4377. **A.** Autoradiogram (Autorad.) and corresponding SDS-PAGE of the autophosphorylation and phosphotransfer reaction employing PK and PKH_497_QR1. Incubation of [γ-^32^P]-ATP radiolabeled truncated version PK with variant PKH_497_QR1 in equal molar amounts. PK, PK+PKR1, and PKH_497_QR1 autophosphorylation reaction served as reference. **B.** Autoradiogram and corresponding SDS-PAGE of the autophosphorylation and phosphotransfer reaction employing PK and PKH_497_QR1D_818_NR2 in equal molar amounts. Incubation of [γ-^32^P]-ATP radiolabeled truncated version PK with non-radiolabeled variant PKH_497_QR1D_818_NR2. PK, PK+PKR1R2, and PKH_497_QR1D_818_NR2 autophosphorylation reaction served as reference. All reactions were stopped after different time points and 10 μl sample resolved on SDS-PAGE, subjected to autoradiography (Autorad.) and Coomassie blue staining to show protein loading (SDS-PAGE).

The purified truncated variant PK was radiolabeled and excess [γ-^32^P]-ATP removed by illustra™ MicroSpin™ G-50 columns (VWR), before PKR1 and PKH_497_QR1 (Fig. 7A) or PKR1R2 and PKH_497_QR1D_818_NR2 (Fig. 7B) were added. Interestingly, no intramolecular phosphotransfer from the kinase PK to one of the fused receiver domains within the variants PKH_497_QR1 or PKH_497_QR1D_818_NR2 was observed (Fig. 7). Both unmodified variants, PKR1 and PKR1R2 were used as reference but still showed autophosphorylation activity due to residual free [γ-^32^P]-ATP (see Fig. S3 for assessing residual free [γ-^32^P]-ATP in the phosphorylated PK sample).

The rather unexpected observation of the lack of intramolecular transphosphorylation prompted us to search for a potential phosphate acceptor. While our results do not exclude a potential function of the intrinsic receiver domains, our study here focused on the identification of the succeeding receiver.

### MA4377 transphosphorylates the downstream encoded single receiver MA4376

As the signal is not transferred from His_497_ to one of the conserved aspartate residues of the fused receiver domains, we were interested in determining whether the downstream encoded single receiver (R) MA4376 may serve as a possible phosphor-acceptor (for genomic localization see Fig. 2).

To examine the intermolecular phosphotransfer of MA4377 to MA4376, the purified recombinant receiver was added to the autophosphorylated kinase without excessive [γ-^32^P]-ATP. Immediately after incubation, a transfer of the radioactive signal to the receiver was observed, while the receiver itself showed no autophosphorylation activity (Fig. 8A). To verify the His1-Asp1 phosphorelay of MA4377 to MA4376, we went ahead to prove that the phosphoryl group is transferred to an aspartate residue within the receiver. Using the variant MA4376D_54_N (RD_54_N), transphosphorylation assays were performed. No signal of RD_54_N was detected after adding the phosphorylated kinase PK which remained phosphorylated, indicating that the phosphoryl group is not accepted by the receiver variant. In addition, these data also verify Asp_54_ of MA4376 as the phosphate acceptor site (Asp1).

**Figure 8.**
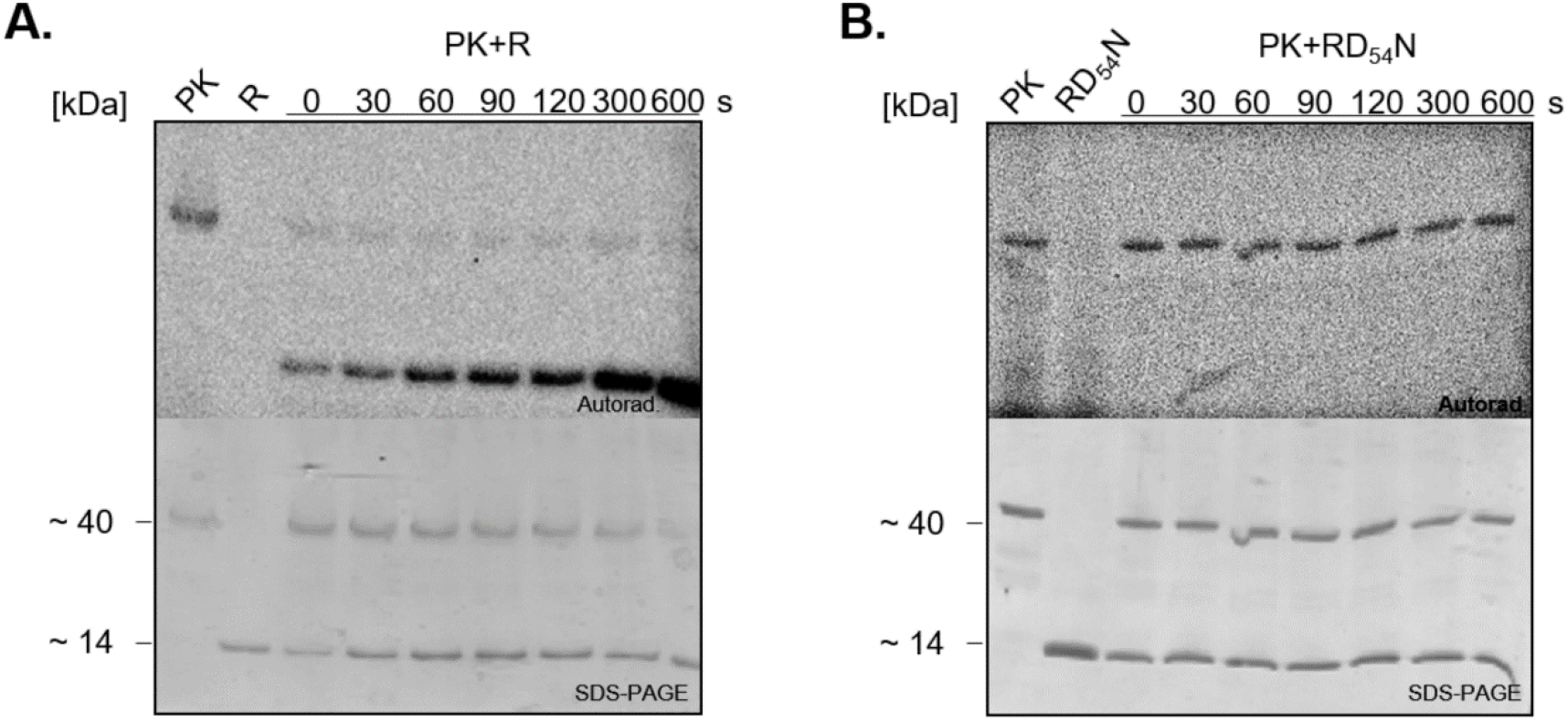
Transphosphorylation assay from MA4377 to MA4376 (R) and the receiver variant MA4376D_54_N (RD_54_N). **A.** Phosphotransfer was assayed by addition of His-tagged R (30 μM) to the autophosphorylated truncated kinase PK (10 μM) in equal molar amounts as reference, PK and R were autophosphorylated alone (10 min). **B.** Phosphotransfer was assayed by the addition of His-tagged RD_54_N to autophosphorylated truncated kinase PK (10 μM). As reference PK and RD_54_N were autophosphorylated alone (10 min). Reactions were stopped after different time points (30-600 s) using 4x SDS sample buffer. Samples (10 μl) were resolved on SDS-PAGE, subjected to autoradiography (Autorad.) and Coomassie blue staining to show protein loading (SDS-PAGE).

In summary, these data show signal transduction from His_497_ of the kinase MA4377 to Asp_54_ of the receiver MA4376 (Fig. 8B).

### A transcription factor as the final phosphocomponent in the phosphorelay of MA4377?

Single receiver domains of bacteria often rely on protein-protein interaction in order to fulfill their function (25). If fused to DNA binding domains, they are able to regulate transcription (26). Thus far, not much is known about the function of bacterial-type single receiver domains in Archaea. Yet, interaction with another protein partner is conceivable. Therefore, we had another closer look at the genomic localization of the MCS. Downstream of MA4377 and MA4376, the transcription factor (TF) MA4375 is encoded (Fig. 2). MA4375 belongs to the Msr family of archaeal transcription factors and related ones have been identified to be involved in the transcriptional activation of corrinoid-methyltransferase fusion proteins (27). MA4375 has the same structure as the above-mentioned TFs, consisting of a helix-turn-helix (HTH) domain and a domain of unknown function 1724 (DUF1724) (27). We therefore asked whether there might be a link between our MCS and the TF MA4375, maybe through transphosphorylation.

The TF itself showed no autophosphorylation activity when incubated with [γ-^32^P]-ATP. Also, no transphosphorylation employing radiolabeled PK and the purified transcription factor was observed (Fig. S4). Possibly, phosphorylation of the TF is achieved through the single receiver MA4376 (for model see Fig.9). For that reason, we tested, whether MA4376 serves as a phosphointermediate in the signal transduction cascade.

**Figure 9.**
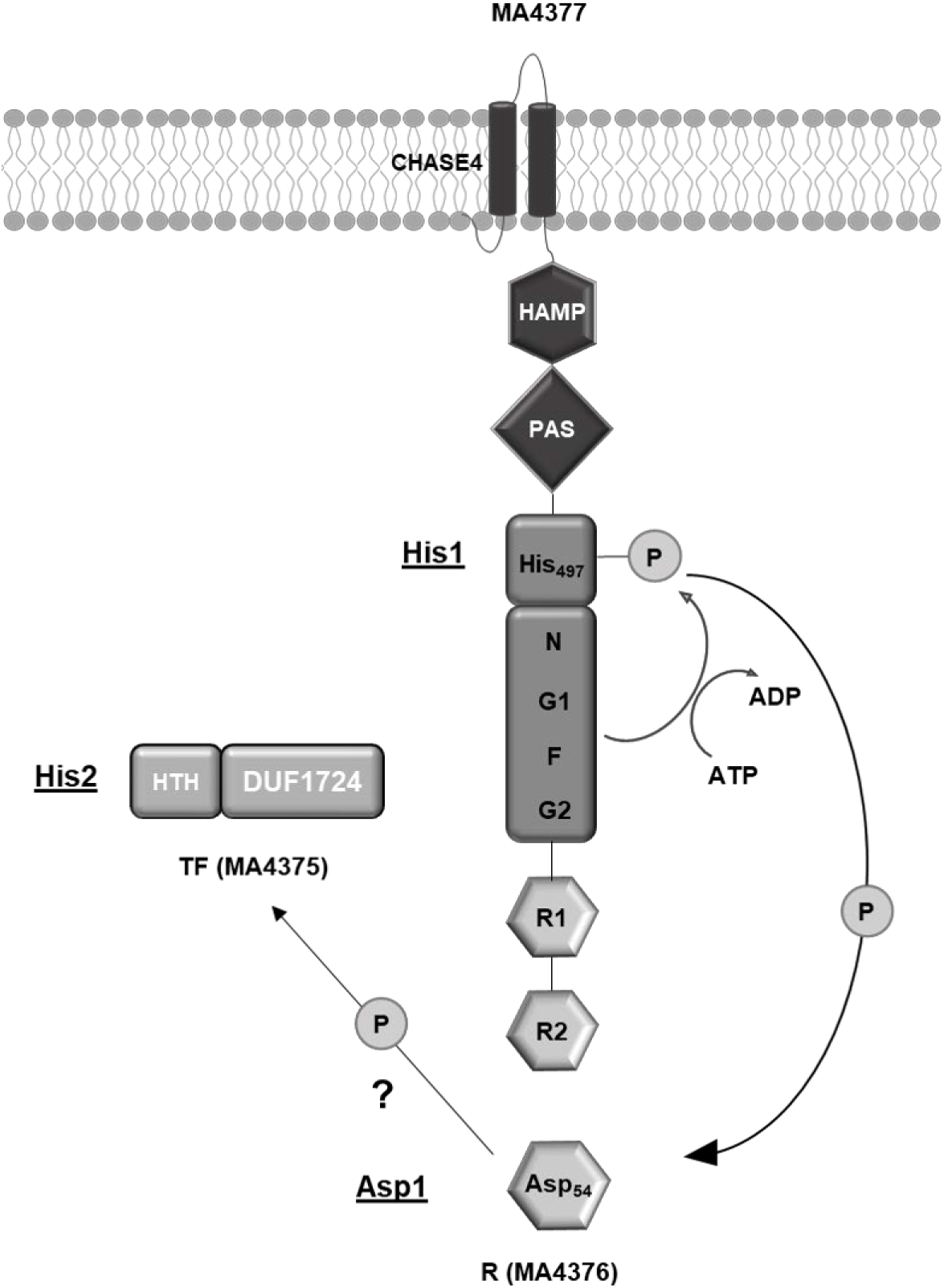
Model of the proposed phosphorelay of MA4377. The kinase MA4377 is as first compound (His1) of the phosphorelay, autophosphorylated at His_497_. The phosphoryl group gets transferred to Asp54 in R (MA4376) which resembles the second compound (Asp2). TF (MA4375) resembles the potential His2 as the next and likely final phosphate acceptor in the phosphorelay.

To investigate this proposed phosphorelay, phosphorylated truncated kinase PK was incubated with the non-radiolabeled receiver MA4376. Subsequently, the transcription factor was added in equal molar amounts and a weak radioactive signal at the size of the TF was detected. The signal intensity of PK attenuated over time as well as the signal of the TF and the strong phosphorylated single receiver R (Fig. 10).

**Figure 10.**
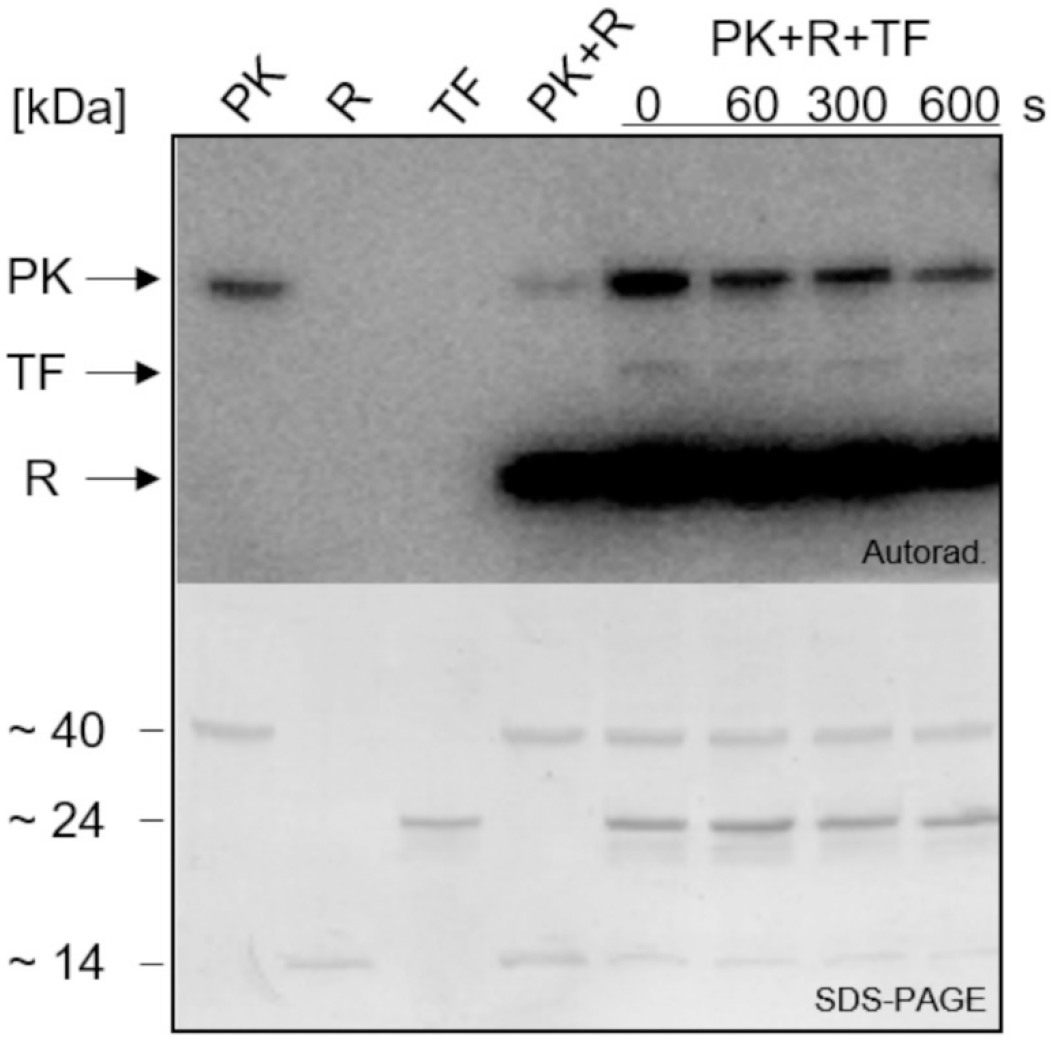
Intermolecular phosphotransfer of MA4377 phosphorelay including receiver (R) and transcription factor (TF). Phosphorelay from PK to receiver MA4376 (R) was assayed and the reaction stopped after 10 min. Truncated version PK (~40 kDa; 9.64 μg) was autophosphorylated for 10 min followed by addition of receiver R (~14 kDa) for another 10 min. Transcription factor (~24 kDa) was added and reaction was stopped after different time points. Samples (10 μl) and autophosphorylation controls (10 μl) were resolved on SDS-PAGE, subjected to autoradiography (Autorad.) and Coomassie blue staining to show protein amount (SDS-PAGE).

The slight phosphorylation of TF points towards an involvement in the phosphorelay of the kinase MA4377.

### Central role of the receiver MA4376 in the phosphorelay of the kinase MA4377

In signal transduction systems single receiver proteins can perceive signals from reverse phosphorelay or cross-talk and transfer it back to the kinase (28, 29). To determine whether MA4376 is able to phosphorylate the kinase, the single radiolabeled R (R-P) was incubated with MA4377. Therefore, R-P had to be obtained, which was implemented by a phosphotransfer from PK to R and subsequent purification of R-P by affinity chromatography (Fig. 11A). In the elution fraction (E) not only the receiver but also the kinase was still present suggesting a tight interaction of both proteins.

**Figure 11.**
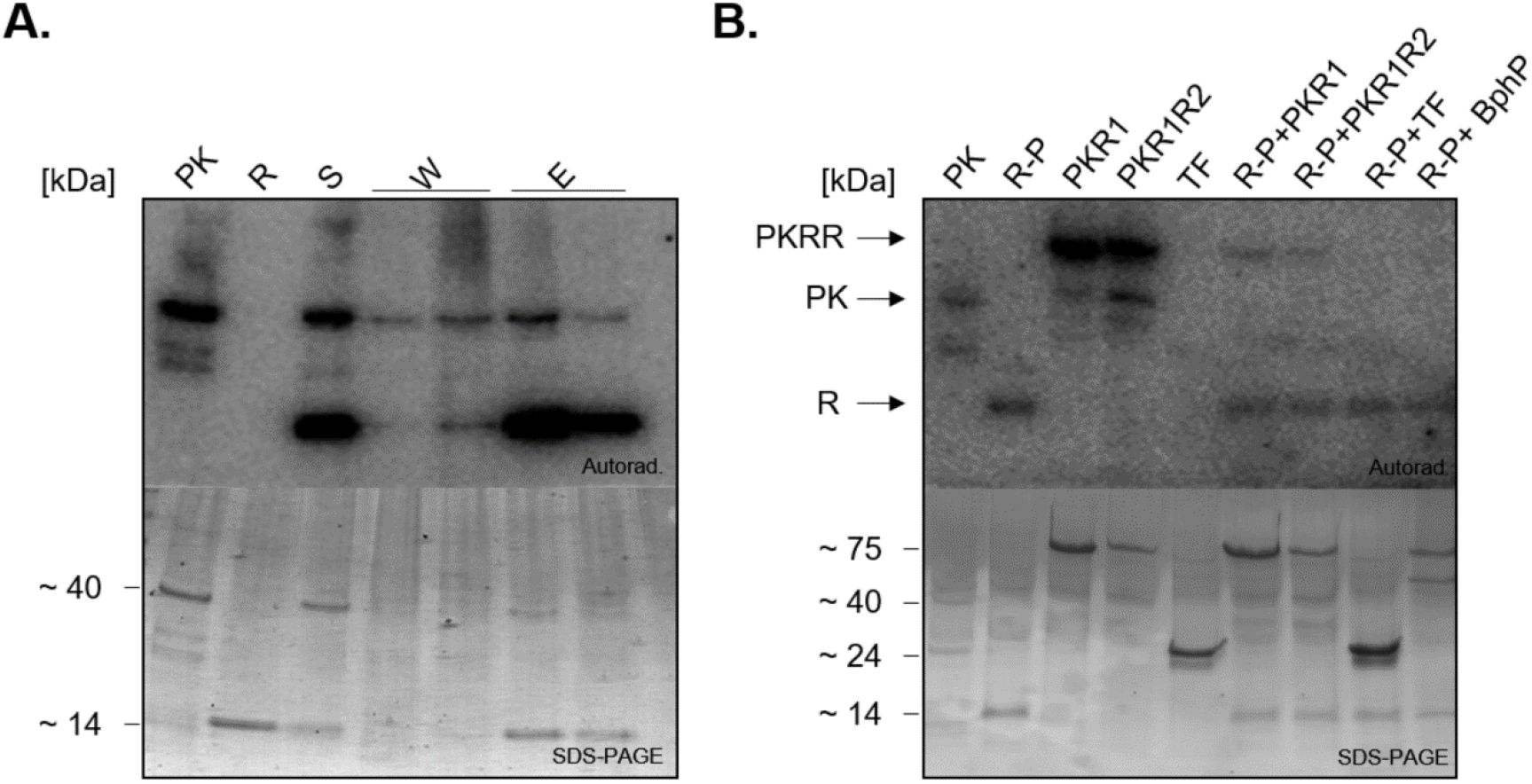
Radioactive His-tagged protein purification of the receiver (R) and reverse phosphotransfer of radiolabeled receiver R to histidine kinase MA4377 and TF MA4375. **A.** Incubation of [γ-^32^P]-ATP radiolabeled truncated version PK with non-radiolabeled receiver MA4376 (R). Supernatant (S), washing (W), and elution (E) fractions of purification steps were resolved on SDS-PAGE and subjected to autoradiography (Autorad.) with subsequent Coomassie blue staining to show protein loading (SDS-PAGE). **B.** Via His-tag affinity chromatography purified receiver (R-P) was incubated with non-radiolabeled truncated versions PKR1 (R-P+PKR1) and PKR1R2 (R-P+ PKR1R2) of MA4377, transcription factor (TF) (R-P+TF) and as control with histidine sensor kinase BphP (R-P+BphP). As a reference autophosphorylation activity of PK, R-P, PKR1, PKR1R2, TF is shown. Samples (10 μl) were resolved on SDS-PAGE, subjected to autoradiography (Autorad.) and Coomassie blue staining to show protein loading (SDS-PAGE).

Phosphorylated receiver was obtained as described in Material and Methods. After desalting, the sole phosphorylated R-P was available, still showing a phosphorylation signal (Fig. 11B). To investigate the reverse phosphotransfer from R-P to the kinase MA4377, R-P was incubated with the truncated versions PKR1 and PKR1R2. Interestingly, our data show a reverse phosphotransfer from R-P to the truncated versions of MA4377 (Fig. 11B).

Having the phosphorylated and purified single receiver available, we furthermore tested whether the receiver by itself is able to phosphorylate the TF or whether the observed phosphotransfer shown in Fig. 10 requires the presence of the kinase. Therefore, the phosphorylated receiver (R-P) was incubated with the TF MA4375. However, no phosphorylation of the TF was observed indicating the importance of the kinase in the phosphorelay. As a reference, the histidine kinase BphP from *Pseudomonas aeruginosa* (30) was incubated with R-P which showed no reverse phosphotransfer (Fig. 11B).

## Discussion

Bacterial signal transduction, whether via a two- or multi-component system involving a phosphorelay, is a well investigated and understood mechanism used by microorganisms in order to respond to environmental changes (4, 6–8). However, their function in Archaea is still largely unexplored, owing in part to only a limited number of genetically accessible organisms being available (31). In screening the ever increasing number of genomes being released, it becomes obvious that these signal transduction systems are also present in Archaea, having possibly radiated from Bacteria via horizontal gene transfer (32). Here, the major question arises as to how these systems have adapted to work together with the archaeal transcription machinery, which possesses components related to that of Bacteria and Eukarya but still being distinct from either. In fact, the archaeal RNA polymerase structurally and evolutionarily resembles eukaryotic RNA polymerase II (33, 34). This study made progress in investigating and understanding of a bacterial- type phosphorelay system in a methanogenic archaeon *M. acetivorans* (3). Using *in vitro* phosphorylation experiments with recombinant protein, we were able to show the autokinase activity of the hybrid kinase MA4377. Although the two fused receiver domains at the C-terminus of the MA4377 protein are not directly involved in the phosphorelay, transphosphorylation to the vicinal encoded single receiver protein, MA4376, was observed. The overlap of 29 nt between both open reading frames might suggest that at some point in their evolution, MA4377 and MA4376 were a single ORF encoding a single hybrid kinase with three R domains. The hybrid kinase RodK from *Myxococcus xanthus* represents such a system, carrying three R domains (35). Only the third R domain is served by an intramolecular phosphotransfer, while the other two are proposed to function as phosphate sinks to modulate the rate of kinase activity inside RodK or to alternative RR in response to incoming signals. Although the authors did not find phosphorylation of R1 and R2 of the *Myxococcus xanthus* kinase, they provide genetic evidence for the distinct function of all three R domains within RodK, with phosphorylation by other kinases being one possible scenario. In the MA4377 and MA4376 system investigated in this study, no phosphorylation of R1 and R2 of the kinase MA4377 was observed. Only transphosphorylation to the single receiver protein MA4376 was detected. Based on the results generated in this study, we propose that MA4376 serves as a central hub to direct the signal into different directions. While it itself is phosphorylated by the kinase, it furthermore supports the phosphotransfer from the kinase to the TF MA4375 (Fig. 10). In addition, it is also able to direct the phosphorylation signal back to the kinase. As all results obtained in this study were from *in vitro* experiments in the absence of most signal input domains (PAS domain still present in employed PK construct), it is necessary to include the membrane-bound signaling domains in future studies to confirm the function of MA4377 *in vivo*. The major signal input domain in MA4377 is likely to be the membrane-bound CHASE domain (36) Although it is widespread in prokaryotes, lower eukaryotes and plants, not much is known about its function. It is predicted to bind various low molecular weight ligands. In the case of MA4377, the CHASE domain might be involved in sensing methylated compounds as alternative substrates for methanogenesis. Data from the Metcalf group support such an assumption (37). This group demonstrated that deletion of the kinase MA4377 and the TF MA4375 in *M. acetivorans* had a significant impact on the transcription of the corrinoid-methyltransferase gene *mtsF* (MA4384), with transcription being upregulated when cells were grown on the alternative substrate trimethylamine (TMA). The synthesis of MtsF was furthermore influenced by the redox-responsive kinase, MsmS (17), indicating that MA4377-MA4376-MA4375 and MsmS might belong to one large signal transduction network. This system furthermore also includes RdmS as a signal input kinase in *M. acetivorans* (Fig. 12), as its deletion had an impact on the transcription of *mtsD* when cells are grown in methanol (18, 37) Interestingly, the kinases MsmS and RdmS are not bacterial-type histidine-kinases and their downstream signal transduction sequence is still not yet fully understood (18).

**Figure 12.**
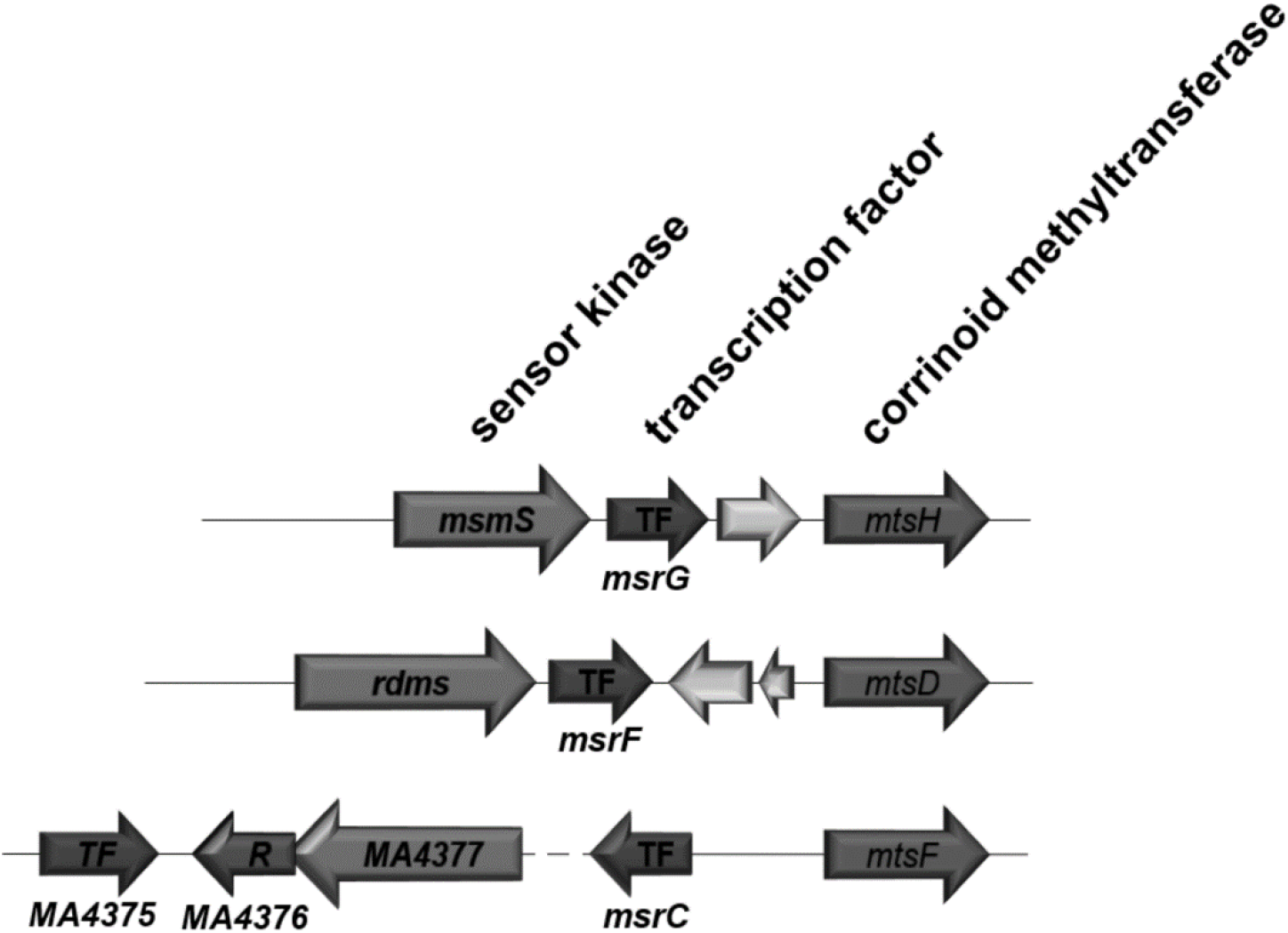
Schematic overview of the methanogenesis related signal transduction system mentioned in the discussion. The genetic organization of kinases, TF and corrinoid methyltransferases are shown as arrows indicating the direction of transcription.

More experiments are needed to clarify whether the receiver MA4376 serves as a central hub in connecting the kinases MsmS, RdmS and MA4377 with the respective TF MA4375, MsrC, MsrF and MsrG, thereby regulating methylotrophic methanogenesis. Whether this interaction involves functional phosphotransfer remains to be determined.

## Material and Methods

### BLASTp and alignment analysis

For Blast analysis BLASTp from NCBI was performed using the non-redundant protein sequence (nr) with an Expect (E) value of 0.05. For detailed analysis special organisms were chosen for BLASTp.

Additional BLASTp analysis were performed with hhpred (MPI Bioinformatics Toolkit) for homology detection and structure prediction of the protein sequence using a E-value cut-off of 1e-3.

For protein structure prediction, the Phyre2 server was used.

Protein sequences used for alignments were searched on NCBI under the domain ‘protein’. The protein sequences were copied as FASTA file into the multiple-sequence-tool Clustal omega (EMBL-EBI) for alignment analysis. No extra settings were executed. The analyses were performed for identifying conserved domains instead of ancestral linkage.

### Construction of expression plasmids

All genes used were PCR amplified from genomic DNA of *M. acetivorans* C2A and cloned according to standard procedures. The constructs from MA4377 were cloned into a *tet* promoter-driven expression system (IBA) with a C-terminal fused StrepII-tag. MA4376 was cloned into the pACYC DUET-1 expression system (Merck) with a C-terminal His-tag. For the transcription factor gene, MA4375, the pET21a (+) expression system with a C-terminal His -tag was chosen as cloning vector. All generated plasmids were verified by Sanger sequencing (SEQ-IT, Kaiserslautern or GATC, Konstanz). All bacterial strains (Table S1), plasmids (Table S2) and oligonucleotides (Table S3) used in this study can be found in the Supplementary data.

### Site directed mutagenesis

Amino acid variants were generated by QuikChange™ site-directed mutagenesis with Pfu-Polymerase and using mismatched primer pairs and the corresponding plasmid template (Table S3).

### Recombinant protein production in *E. coli* BL21(DE3)

Cultures were grown at 37 °C in LB-HS medium to an OD_578_ of ~0.5 and gently cooled down to 17 °C. Expression of MA4375 pET21a (+) was induced with 0.5 mM isopropyl β-thiogalactoside (IPTG). *Tet* promoter driven systems were induced with 200 ng/μl anhydrotetracycline (AHT). Cultures of either promoter system were further incubated for 16 h at 17 °C and harvested for 10 min at 3500 x g. The cell pellet was stored at −20°C or directly used for protein purification.

### Purification of recombinant produced StrepII-tagged proteins

*E. coli* cell pellets containing StrepII-tagged proteins were resuspended in freshly prepared buffer W (100 mM Tris/HCl, pH 8.0, 300 mM NaCl, 1 mM EDTA) and freshly added 0.25 mM 4-(2-aminoethyl) benzene-sulfonyl fluoride and 1 mM 1,4-dithiothreitol (DTT). In order to purify the StrepII-tagged proteins, affinity-chromatography using a Strep-Tactin® gravity flow column (IBA) was used. A column containing Strep-Tactin was equilibrated with buffer W. The filtered lysate was added onto the column and the flow through was collected. To remove all remaining lysate debris the column was washed with buffer W (10 CV). For elution, buffer E (buffer W containing 2.5 mM D-desthiobiotin) was used. Purified protein was dialyzed against Tris buffer pH 8.0 (50 mM Tris, 150 mM NaCl, 5 mM MgCl_2_, 50 mM KCl) and concentrated using Amicon concentrator devices (molecular weight cut-off, 10 000 and for full-length protein 50 000; Merck Millipore). Recombinant His-tagged proteins (MA4376, MA4376D_54_N and MA4375) were purified with the ÄKTAprime plus system using HisTrap HP (GE healthcare) columns. The cell pellets were resuspended in buffer W_His_ (buffer W with additional 10 mM imidazole) and loaded onto the HisTrap HP column which was previously equilibrated with buffer W_His_. His-tagged proteins were eluted using W_His_ buffer containing 150 mM imidazole.

### Size exclusion chromatography

Size exclusion chromatography was performed on a Superose™ 6 10/300 GL column (GE Healthcare) using 50 mM NaPOH_4_ buffer at pH 7.0.

As standard proteins Apoferritin (MW = 443 kDa), ß-Amylase (MW = 200 kDa), alcohol dehydrogenase (MW = 150 kDa), Albumin (MW = 66 kDa) and carbonic anhydrase (MW = 29 kDa) were used. The void volume was determined using Blue Dextran (2000 kDa).

### Radioactive *in vitro* autophosphorylation and phosphotransfer assays

In order to investigate the autophosphorylation activity of the desired protein or protein variant a reaction mix of 10 μM of the purified protein, 2.5 μl of 5xTris-buffer pH 8.0 (50 mM Tris, 150 mM NaCl, 5 mM MgCl_2_, 50 mM KCl) and ad. H_2_O up to a total volume of 10 μl was combined in a sealed short thread vial (VWR). For reducing condition freshly prepared DTT was added to a final concentration of 2 mM. The reaction mix was flushed with N_2_ using a cannula (Braun) for 15 sec whereas air bubbles were removed to create a reduced environment in the vial. To start the reaction 2.5 μl of the ATP Mix (0.25 μl ATP (10 mM), 0.25 μl [γ-^32^P]-ATP (1850 kBq), 2 μl H_2_O) was added and samples were taken after different time points, and the reaction was stopped by the addition of 4x SDS sample buffer.

To investigate the phosphorelay inside the kinase MA4377, 10 μM of PK were autophosphorylated for 10 min. After removing excessive ATP with illustra™ MicroSpin™ G-25 Columns (GE healthcare), the protein was added to an extra vial containing 10 μM of the phosphate acceptor constructs (PKH_497_QR1, PKH_497_QR1D_818_NR2, R, RD_54_N and TF). All phosphate acceptor constructs were reduced with fresh DTT (2 mM) and flushed with N_2_ before phosphotransfer reaction.

To separate the phosphate acceptor construct from PK and to remove imidazole in the case of phosphorylated R, subsequent to affinity chromatography, Sephadex G-25 (GE) column were used.

All taken samples (10 μl) were quenched with 4x SDS sample buffer 100 mM Tris pH 8.8 (HCl), 110 mM SDS, 40 % (w/v) glycerin, 3 mM bromophenol blue, 2 mM β-mercaptoethanolethanol) and then loaded without prior heating on a 4-20% gradient polyacrylamide gel and separated by denaturing sodium dodecyl sulfate-polyacrylamide gel electrophoresis (SDS-PAGE) (BioRad). The gel was exposed to a PhosphorImager screen in a cassette containing an enhancing screen, overnight at room temperature. The signals were recorded using a PhosphorImager (GE Healthcare Perkin Elmer Cyclone Typhoon FLA 7000). To visualize the proteins, the same gel was stained in Coomassie Brilliant Blue G250, destained in 30 % (v/v) ethanol, 10 % (v/v) acetic acid and documented by scanning with HP Scanjet 300 (HP).

To quantify the intensity of the increasing autophosphorylation activity of PK (10 μM) ImageJ (64-bit Java 1.8.0_172) Color_Pixel_Counter was used. The sample taken after 600 s referred as reference and was valued as 100% of signal intensity.

### Acid–base treatment of phosphorylated protein

The proteins used in this work were phosphorylated, then separated via SDS-PAGE and transferred onto a PVDF membrane (18 min, 15 V) by Trans-Blot^®^ SD semi-dry Western blot (BioRad). The membrane was exposed as described before overnight to a PhosphoImager screen. After the detection of the radioactive signals with a PhosphoImager (GE Healthcare Perkin Elmer Cyclone Typhoon FLA 7000), the membrane was incubated for 1 h at RT in 6 M HCl, 3 M NaOH and, as reference, in H_2_O. Phosphorylation signals were again evaluated by exposing the membrane overnight as described above to a PhosphoImager screen and subsequently detected. To confirm protein transfer, the PVDF membrane was stained with Ponceau S to visualize the proteins.

### Histidine residue modifying reagent diethylpyrocarbonat (DEPC)

By using the reagent diethylpyrocarbonat (DEPC) (Carl Roth), which modifies histidine residues, the phosphorylated amino acid residue of a kinase can be determined (38). The reagent was diluted in EtOH to the required concentration and incubated for 1 h at room temperature with the purified protein in a total volume of 7.5 μl. The sample was subsequently autophosphorylated for 10 min and used in a kinase assay to evaluate if the reagent is masking the histidine residue under investigation.

## Acknowledgements

This work was in part supported by a grant from the Deutsche Forschungsgemeinschaft, to NFD. We thank Loriana Blasius and Nora Georgiev for critically reading of the manuscript and Dr. Michelle Gehringer for English language editing.

